# Expression spectrum of TE-derived transcripts in human adult tissues

**DOI:** 10.1101/2025.03.29.646092

**Authors:** Benpeng Miao, Xinlong Luo, Amina Ademovic, Yushan Yang, Tao P. Wu, Bo A. Zhang

## Abstract

Transposable elements (TEs) are vital components of eukaryotic genomes and have played a critical role in genome evolution. Although most TEs are silenced in the mammalian genome, increasing evidence suggests that certain TEs are actively involved in gene regulation during early developmental stages. However, the extent to which human TEs drive gene transcription in adult tissues remains largely unexplored. In this study, we systematically analyzed 17,329 human transcriptomes to investigate how TEs influence gene transcription across 47 adult tissues. Our findings reveal that TE-derived transcripts are broadly expressed in human tissues, contributing to both housekeeping functions and tissue-specific gene regulation. We identified sex-specific expression of TE-derived transcripts regulated by sex hormones in breast tissue between females and males. Our results demonstrated that TE-derived alternative transcription initiation significantly enhances the variety of translated protein products, e.g., changes in the N-terminal peptide length of WNT2B caused by TE-derived transcription result in isoform-specific subcellular localization. Additionally, we identified 68 human-specific TE-derived transcripts associated with metabolic processes and environmental adaptation. Together, these findings highlight the pivotal evolutionary role of TEs in shaping the human transcriptome, demonstrating how conserved and human-specific TEs contribute to transcriptional and translational innovation in human genome evolution.

## Introduction

Transposable elements (TEs) are mobile genetic sequences, that constitute significant components of animal and plant genomes, have been found as the pivotal driving forces in shaping genome architecture and functio^1-4^. TEs have driven genetic innovation through mechanisms such as promoting genome rearrangements, introducing novel regulatory sequences, and facilitating the emergence of new genes. As important sources of genetic variability, TE insertions in the host genome provided the raw material for both evolutionary adaptation and biological function conformation. For example, TEs contribute significant components of enhancers in the human genome, with strong tissue-specific activation patterns^1,5-7^. In the mouse genome, SINE elements carry nearly 30% of CTCF binding sites and contribute to the evolutional conservation of chromatin topological associated domains^8-11^. Furthermore, specific TEs are reactivated during stress or in particular cell types, such as stem cells^12,13^ and immune cells^14,15^, highlighting their functional diversity in maintaining cellular homeostasis and mediating adaptive responses^16-18^. LINE1 elements are expressed during early embryonic development and regulate pluripotency-associated genes^19,20^. SINE B2 elements were discovered to enhance the transcription of heat shock protein genes under stress^21^, and could work as enhancers in response to IFN stimulus to regulate target genes^22^.

TEs can also function as alternative promoters by providing transcription start sites (TSSs) and regulatory sequences that drive gene transcription in various biological contexts^23-26^. TEs often contain various binding sites of transcription factors and core promoter motifs (e.g., TATA box-like sequences), after integrating near or within genes, TE insertion can provide novel sequences with the ability to recruit transcriptional machinery and drive the transcription elongation. In human placental development, LTR elements from endogenous retroviruses regulate imprinted genes like PEG10 and RTL1, essential for proper placental function^27^. In human airway epithelia cells, the Alu element can transcribe a specific isoform of IL33 and associate with human chronic obstructive pulmonary disease^16^. In mouse embryo development, hundreds of TEs were found to initiate specific gene’s transcription in a tissue-specific fashion with temporal patterns^28^. However, how many human genes can be transcribed from TE-derived TSSs has not been comprehensively studied. Especially in the mature tissues, how the TE-derived TSSs regulate the host gene expression and are further involved in functional maintenance is largely unclear.

In this study, we analyzed the 17,329 human transcriptomes of distinct adult tissue types, generated from GTEx consortium, to systematically evaluate the TE-derived transcripts in 44 female tissues and 43 male tissues. We discovered that TE-derived transcripts are broadly expressed in human tissues, contributing to both housekeeping functions and tissue-specific gene regulation. In the human genome, we identified 3528 protein-coding genes and 944 lincRNA genes that can use TE-derived TSS to initiate transcription. We identified sex-biased expression of TE-derived transcripts in breast tissue between females and males, and more than half of these TEs harbored sex hormone receptor binding motifs, such as AR and ESR1. However, minor expression differences of TE-derived transcripts were observed in other tissues. We further demonstrated that TE-derived alternative transcription initiation enhanced protein translation diversity during the evolution. In our study, 1009 protein-coding genes used TE-derived TSS to transcribe the transcript isoforms that can be further translated to proteins that have altered peptide length, when compared to canonical non-TE-derived TSS. Finally, we examined the conservation of these human TEs that can initiate gene transcriptions among vertebrates, and identified 68 human-specific TEs, that can drive the transcription of genes associated with metabolic processes and environmental adaptation. Collectively, these findings underscore the critical evolutionary impact of transposable elements (TEs) in shaping the human transcriptome, illustrating how both conserved and human-specific TEs drive transcriptional and translational diversity, thereby influencing the evolutionary history of the human genome.

## Result

### TE-derived transcripts in human tissues

To investigate the expression patterns of TE-derived transcripts in human tissues, we analyzed all transcription start sites (TSSs) of annotated transcripts in the human genome using GENCODE v26. We found 21% of human genes can utilize TE-derived TSSs, comprising 5,330 protein-coding genes and 2,673 non-coding genes [Supplementary figure 1a]. To evaluate the expression of these TE-derived transcripts, we examined the GTEx dataset (release V8) and found 4,743 and 4,482 TE-derived transcripts from protein-coding genes expressed (>0.1 RPKM) in male and female tissues, respectively [Figure 1a, supplementary file 1]. On average, each tissue expressed approximately 3000 protein-coding and ∼400 lincRNA TE-derived transcripts. Most of these transcripts were expressed at relatively low levels, with ∼65% of protein-coding and ∼80% of lincRNA TE-derived transcripts exhibiting expression levels below 1 RPKM (Figure 1b and supplementary figure 1b). Interestingly, more expressed lincRNA transcripts were observed in male tissues (1,103) than in female tissues (809). Notably, the testis displayed the highest number of TE-derived lincRNA transcripts, with 856 transcripts identified [Figure 1b].

**Figure 1.**
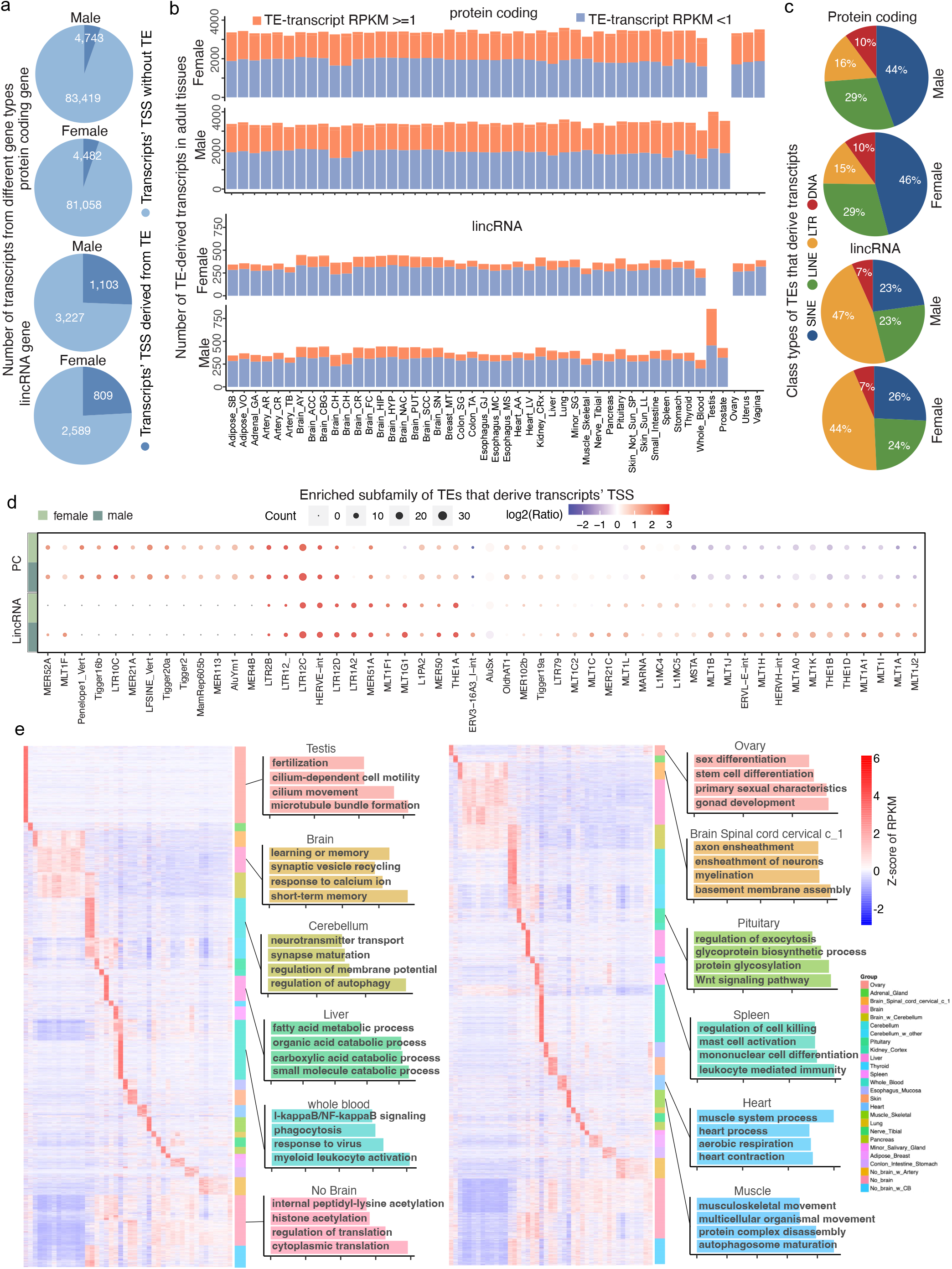
Expression of TE-derived transcripts in human adult tissues. **a)**. The total number of TE-derived transcripts identified in lincRNA and protein-coding genes across male and female tissues. **b)**. Numbers of TE-derived transcripts across different adult tissues in females and males. **c)**. The classes of transposable elements (TEs) derived transcripts of lincRNA and protein-coding genes, with colors representing different TE classes. **d)**. The subfamily enrichment of TEs contributing to transcript TSSs in lincRNA and protein-coding genes is illustrated. The size of each dot represents the number of individual TEs within a subfamily. The ratio indicates the fraction of transcripts derived from specific TE subfamilies relative to their overall fraction in the human genome. PC: protein-coding genes. **e)**. Heatmap showing the Z-score expression of protein-coding TE-derived transcripts across tissues for males (left) and females (right). Color-matched bar plots indicate enriched gene ontology results in different tissues.

Among the TE-derived transcripts, nearly half of the TSSs for protein-coding genes were contributed by SINE elements, while LTR elements accounted for about half of the TSSs for lincRNA genes [Figure 1c]. Several previously reported TE subfamilies, including MER52A, LFSINE, and LTR12C, were highly enriched in TE-derived protein-coding transcripts [Figure 1d, supplementary file 1]. On average, ∼3000 protein-coding and ∼300 lincRNA TE-derived transcripts were expressed in both male and female tissues [supplementary figure 1c], highlighting TE-derived transcription as a prevalent mechanism in the human genome. In addition to commonly expressed TE-derived protein-coding transcripts, we identified tissue-specific TE-derived transcripts in each tissue [Supplementary file 2, supplementary figure 1d-e]. These tissue-specific transcripts were strongly associated with the unique biological functions of the corresponding tissues [Figure 1e]. These findings suggest that TEs have been domesticated as alternative promoters to enhance tissue-specific gene regulation during evolution, reflecting their critical role in shaping the transcriptomic landscape.

### Sex-biased expressed TE-derived transcripts in distinct human tissues

To investigate the sex-biased expression patterns of TE-derived transcripts in human tissues, we conducted differential expression analysis between females and males for each tissue. In most tissues, the majority of TE-derived transcripts exhibited limited sex-biased expression (fold change <1.5). However, approximately 9% of TE-derived protein-coding transcripts and 11% of lincRNA transcripts displayed significant sex-biased expression (fold change >1.5) [Figure 2a, supplementary file 3]. Notably, in breast tissue, we identified 312 protein-coding transcripts and 83 lincRNA transcripts with substantial sex-biased expression between females and males. Most of these transcripts were highly expressed in females [Figure 2b, supplementary file 3], including key genes associated with mammary gland development, such as KRT15, FGGY, and LIMCH1. KRT15, expressed in the basal compartment of the mammary gland, marks long-lived stem cells capable of generating all epithelial components of the anagen follicle^29^. Interestingly, KRT15 is transcribed from a MIRb-derived promoter, and its activation is directly regulated by ESR1 binding in females [Figure 2c]. Another example is ACSM3 (Acyl-CoA Medium-Chain Synthetase-3), which inhibits proliferation, motility, and stem cell properties of breast-invasive carcinoma^30,31^. Two transcription start sites (TSSs) of ACSM3 located in HERV15-int loci are specifically activated by ESR binding in female breast tissues [Figure 2d]. Gene ontology analysis of these sex-biased TE-derived genes revealed enrichment in mammary gland-related functions, including vascular processes, vasculature development, monatomic anion transport, and pathways associated with breast cancer [Figure 2e].

**Figure 2.**
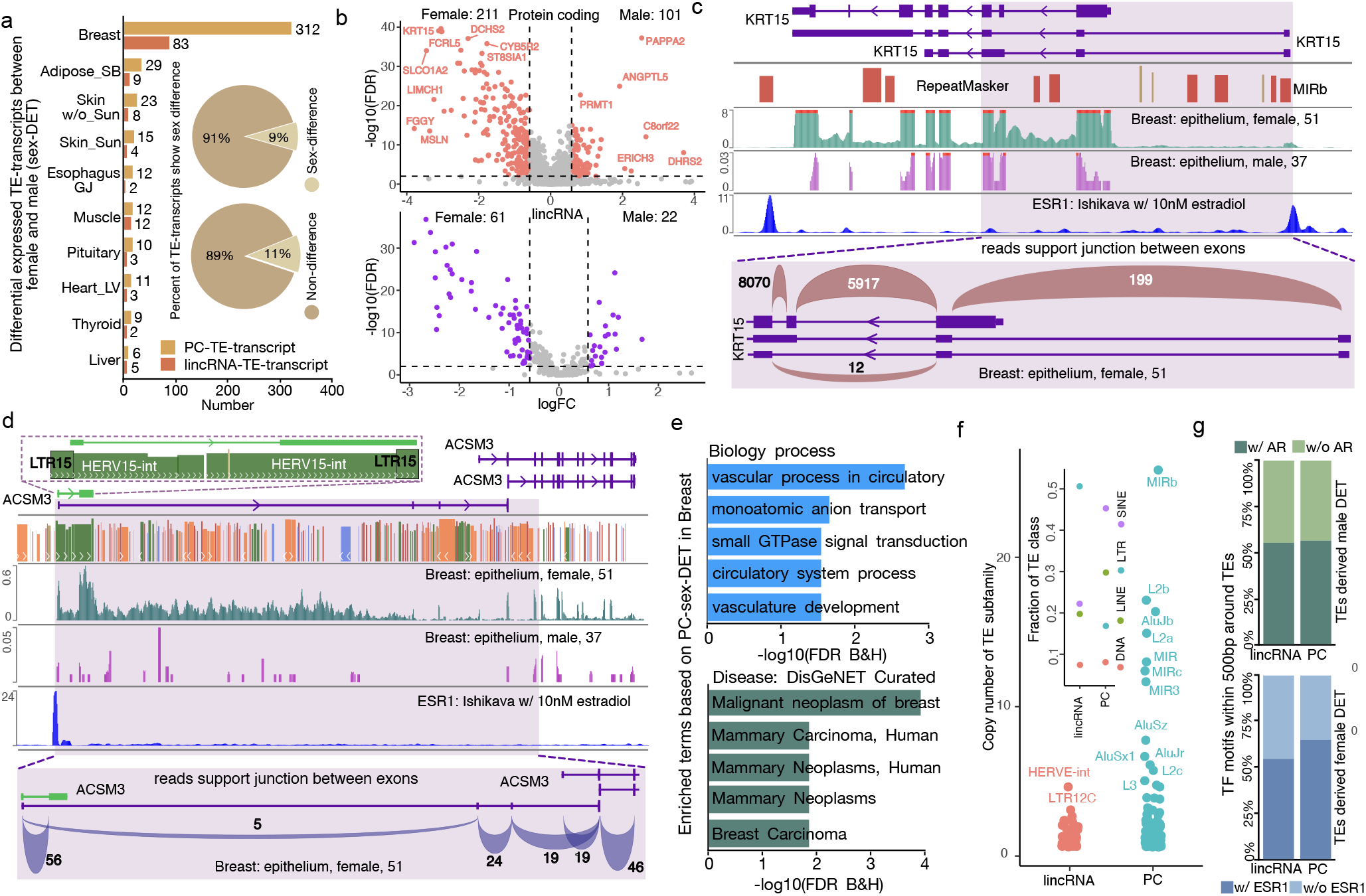
Differentially expressed TE-derived transcripts between males and females. **a)**. Number of differentially expressed TE-derived transcripts (sex-DETs) between males and females across tissues for lincRNA and protein-coding genes. **b)**. The volcano plots display sex-DETs from protein-coding genes (top) and lincRNAs (bottom) in breast tissue. The y-axis represents the -log10 FDR, while the x-axis represents the log2 fold changes of sex-DETs between females and males. **c)**. The MIRb-derived transcript of KRT15 gene. **d)**. The LTR15-derived transcript of the ACSM3 gene is shown. For both **(c)** and **(d)**, the top panel illustrates RNA-seq signals from female and male breast samples, along with ChIP-seq signals for the ESR1 gene. The bottom panel displays reads supporting the exon junctions of transcripts in female breast samples. **e)**. Enriched biological processes and diseases associated with protein-coding genes containing sex-DETs in breast tissue. **f)**. TEs that contribute to sex-DETs in breast tissue for protein-coding genes and lincRNAs. The inner dot plot represents the fraction of TE classes, while the outer dot plot displays the number of individual TEs within each subfamily. **g)**. The percentage of TEs intersecting with AR and ESR1 transcription factor binding motifs. The top panel shows the percentage of TEs driving up-regulated sex-DETs in males that overlap with AR binding motifs, while the bottom panel displays the percentage of TEs driving up-regulated sex-DETs in females that overlap with ESR1 binding motifs.

We further examined the TE subfamily distribution of sex-biased TE-derived promoters. The MIRb and L2b subfamilies significantly contributed to sex-biased protein-coding TE-derived transcripts, while HERV15-int and LTR12C were predominantly associated with lincRNAs in both male and female breast tissues [Figure 2f]. Additionally, certain subfamilies showed distinct sex-biased activity: L2a and AluJb were enriched in females, whereas AluY was exclusively activated in male breast tissue [supplementary figure 2]. Motif analysis revealed a strong enrichment of sex hormone receptor binding sites in domesticated TE promoters. Notably, more than 50% of female-highly expressed genes had ESR1 binding sites in their promoter regions, while approximately 60% of male-highly expressed genes had promoters containing AR binding sites [Figure 2g]. These findings highlight the significant role of TEs in shaping sex-specific gene regulation, particularly through interactions with hormone receptor signaling pathways.

### TE-derived dominant transcriptions contribute to housekeeping functions in the human body

Although thousands of TE-derived transcripts are identified across various tissue types, many are expressed at relatively low levels [Figure 1b]. However, when assessing the contribution of TE-derived transcripts to the overall expression of their host genes, we observed that approximately 30% of these transcripts (1300 in females and 1619 in males) dominantly contribute to more than 50% of their host gene expression in at least one tissue type [Figure 3a, supplementary file 4].

**Figure 3.**
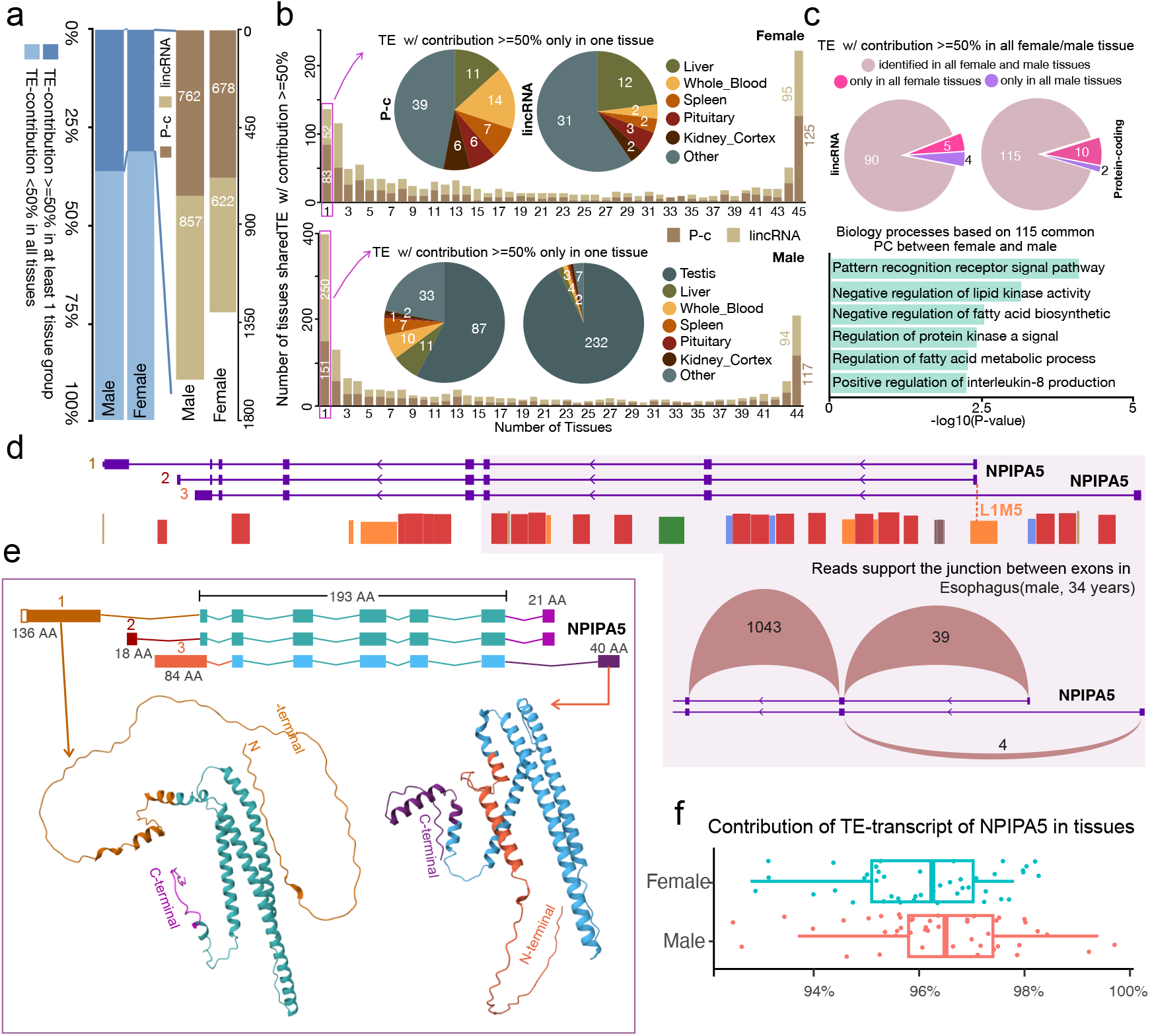
The contribution of TE-derived transcripts in gene expression. **a)**. Number of genes with more than 50% of their expression contributed by TE-derived transcripts in at least one tissue. The left panel shows that approximately 30% of such genes have their expression predominantly driven by TEs, comprising around 800 lincRNA genes and 700 protein-coding genes (right panel). TE-contribution refers to the expression contribution from TE-derived transcripts. P-c denotes protein-coding genes. **b)**. Tissue distribution of lincRNA and protein-coding genes predominantly driven by TEs for females (top) and males (bottom). Bar colors indicate protein-coding genes (P-c) and lincRNAs. Many genes were found in either one tissue or across all tissues. Pie charts show tissue-specific genes, with a notable concentration in the testis. Pie chart colors represent different tissues. **c)**. Sex distribution of protein-coding genes and lincRNAs with more than 50% TE contribution (top) and enriched biological processes based on common protein-coding genes between females and males (bottom). **d)**. L1M5-derived transcripts of the NPIPA5 gene. The bottom-right plot shows reads supporting the junctions between the TE-derived exon and the subsequent exon in a male esophagus sample. **e)**. Protein structures of TE-derived and canonical non-TE transcripts of the NPIPA5 gene. **f)**. Boxplot showing the expression contribution of TE-derived transcripts in the NPIPA5 gene across tissues, with each dot representing the contribution in a single tissue.

Interestingly, the tissue specificity of these dominant TE-derived transcripts exhibits a bimodal distribution. 220 TE-derived transcripts are expressed across all tissue types in female tissues, while 135 are restricted to a single tissue. Notably, 23 single-tissue-expressed TE-derived transcripts were identified in the liver, suggesting a role in female-specific metabolic processes. In males, 401 TE-derived transcripts are expressed exclusively in a single tissue, with 319 specifically expressed in the testis. This highlights the unique epigenetic and transcriptomic landscape of male germ-line cells [Fig. 3b, supplementary figure 3]. Additionally, 115 protein-coding and 90 lincRNA TE-derived transcripts were ubiquitously expressed in all tissues in both males and females. These transcripts, associated with housekeeping functions, are highly enriched in core biological processes such as receptor signaling pathways, lipid metabolism, and interferon regulation [Figure 3c]. This evidence suggests that TEs were co-opted during early human evolution to serve as promoters for initiating housekeeping gene expression critical to fundamental biological functions.

An example of such functional adaptation is NPIPA5, a member of the nuclear pore complex-interacting protein family, which plays a critical role in regulating the mitotic cell cycle by influencing centrosome segregation, centriole maturation, and spindle orientation—processes essential for proper cell division^32^. In human tissues, three major isoforms of NPIPA5 have been identified, two of which are transcribed from an intergenic L1M5-derived transcription start site (TSS) located within the first intron [Figure 3d]. Compared to the canonical isoform, this primate-specific L1M5-derived TSS produces two alternative isoforms with a novel 21-amino-acid N-terminal peptide sequence and varying lengths of C-terminal peptides [Figure 3e]. Remarkably, the L1M5-derived TSS accounts for over 90% of NPIPA5 transcription in human tissues [Figure 3d, 3f]. These findings underscore the critical role of transposable element (TE) insertions in generating novel protein products and reshaping human gene regulation, illustrating how TEs contribute to evolutionary innovation.

### TE-derived transcripts increase the diversity of protein isoforms

Transposable element (TE)-derived alternative transcription initiation sites have the potential to alter protein products by generating transcript diversity [Figure 3d]. To systematically assess such diversity induced by transposable elements, we classified TE-derived transcripts of protein-coding genes into distinct categories based on GENCODE and Ensembl annotations (see Methods). Our analysis revealed that nearly 60% of these transcripts are non-translatable, encompassing retained-intron transcripts, processed transcripts, and nonsense-mediated decay transcripts. The remaining ∼40% (1759 transcripts) are annotated to encode protein products [Figure 4a]. Among these categories, we observed a distinct enrichment of short interspersed nuclear elements (SINEs) in retained-intron transcripts, accounting for 53% of individual copies within this group [Figure 4b]. This suggests that SINE-derived promoters may influence abnormal splicing mechanisms during transcription elongation. Gene ontology (GO) analysis further demonstrated functional differentiation among transcript categories. TE-derived protein-coding transcripts were enriched in pathways related to inflammatory responses, PLC signaling, and TGF signaling. In contrast, TE-derived retained-intron transcripts showed enrichment in protein modification and chromatin remodeling pathways [Figure 4c, supplementary figure 5].

**Figure 4.**
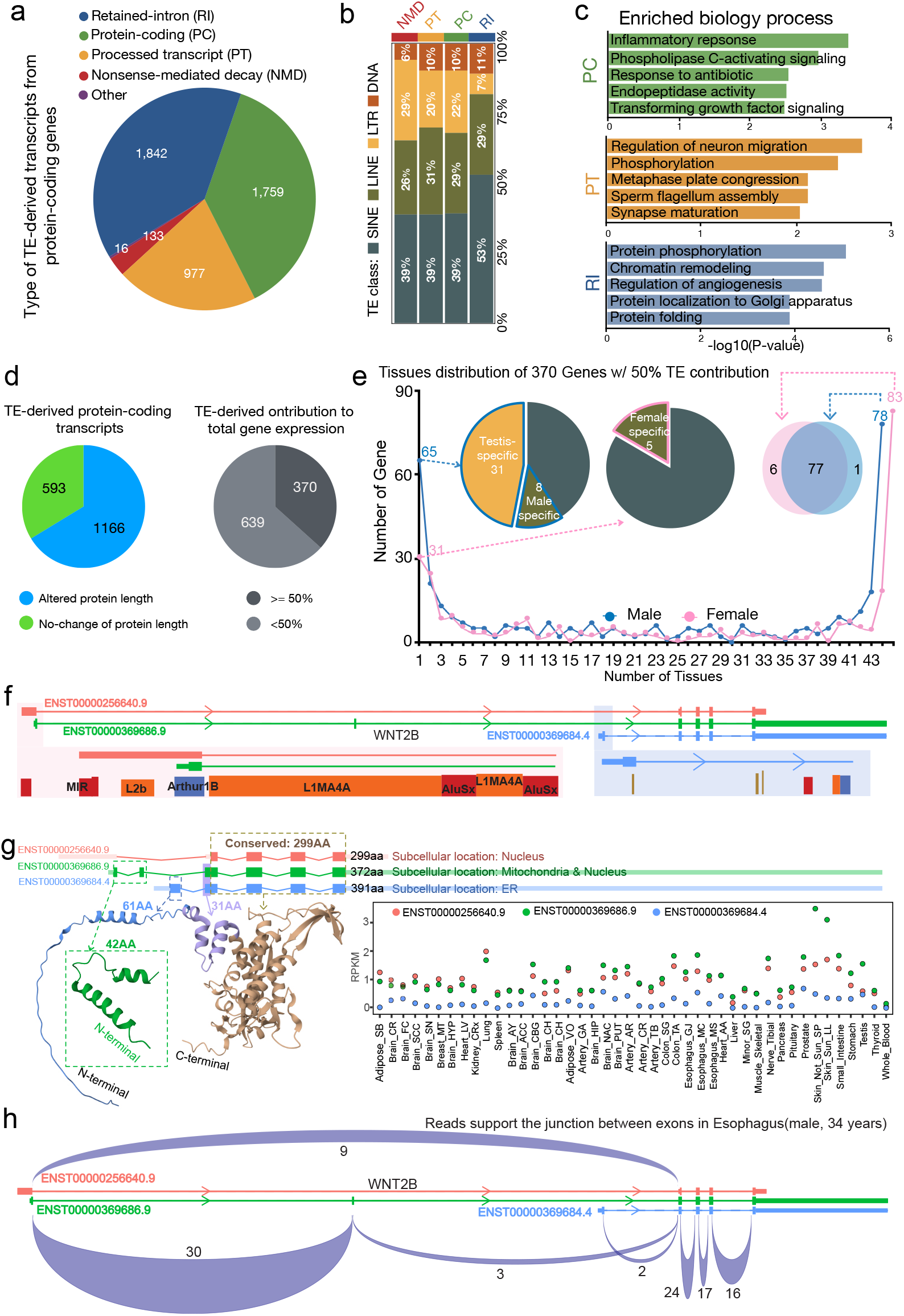
TE-derived protein-coding transcripts contribute to enhancing protein diversity. **a)**. Different types of TE-derived transcripts from protein-coding genes include retained-intron (RI), protein-coding (PC), processed transcripts (PT), and nonsense-mediated decay (NMD) transcripts, which constitute the major fractions of TE-derived transcripts. **b)**. The classes of TEs contributing to different types of transcripts. Approximately 50% of retained-intron transcripts’ TSSs are derived from SINE elements. **c)**. The enriched biological processes associated with different transcript types. **d)**. Protein length and expression contribution of TE-derived protein-coding transcripts. Among these, 1,166 (∼66%) TE-derived protein-coding transcripts from 1,009 genes exhibited altered protein lengths compared to non-TE protein-coding transcripts. Additionally, 379 genes had more than 50% of their expression contributed by these TE-derived protein-coding transcripts. **e)**. Tissue distribution of 370 protein-coding genes with more than 50% TE contribution. Pink and blue lines represent females and males, respectively. The two pie charts on the left show the sex-specific distribution of genes identified in only one tissue for males and females. **f)**. Three transcripts of the WNT2B gene include the MIR and Arthur1b-derived transcripts (ENST00000256640.9 and ENST00000369686.9) and a non-TE-derived transcript (ENST00000369684.4). **g)**. The protein production and subcellular localization of the three WNT2B gene transcripts are shown. The dot plot on the right displays the RPKM values of different transcripts across tissues, with TE-derived transcripts being predominantly expressed in most tissues. **h)**. The sashimi plot of the male esophagus sample illustrates reads supporting the junctions of exons in both TE-derived and canonical non-TE transcripts of the WNT2B gene.

**Figure 5.**
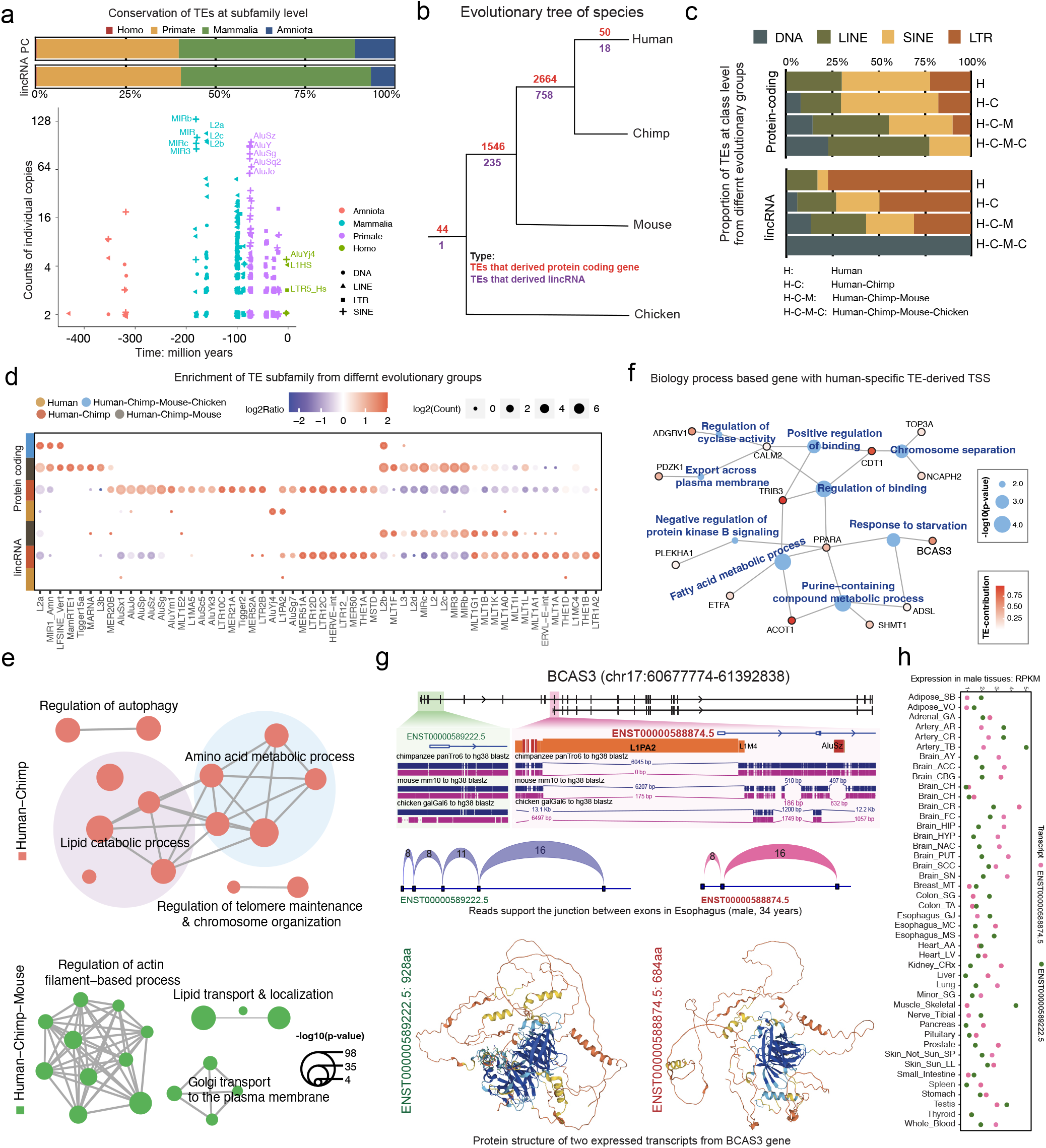
The evolutionary conservation of TEs that derived transcripts of lincRNA and protein-coding genes. **a)**. The evolutionary conservation of TEs contributes to transcripts at the subfamily level. The top panel illustrates the taxonomic groups of TEs that derived transcripts, with colors representing different taxa groups. The bottom plot displays the estimated evolutionary age (in million years) of the TEs, with shapes representing different TE classes. **b)**. An evolutionary tree of TEs based on sequence conservation across human, chimpanzee, mouse, and chicken genomes. Different colors indicate TEs that derived TSSs from protein-coding genes or lincRNAs. **c)**. The class proportion of TEs from different evolutionary groups for protein-coding genes and lincRNAs is displayed. For protein-coding genes, SINE elements occupy a high fraction in H and H-C groups, whereas LTR elements dominate for lincRNAs. **d)**. Subfamily enrichment of TEs from different evolutionary groups for protein-coding genes and lincRNAs. The size of each dot represents the number of individual TEs within a subfamily across different evolutionary groups. **e)**. Enriched biological processes based on genes with transcripts derived from TEs in different evolutionary groups. Red indicates processes enriched in the Human-Chimp group, while green represents processes enriched in the Human-Chimp-Mouse group. Connected lines indicate shared genes between different biological processes. **f)**. Enriched biological processes based on genes with transcripts derived from human-specific TEs. Blue dots represent biological processes, with size indicating the p-value. Black circles represent genes, with red color indicating the level of TE contribution to the gene. Grey lines connect genes to their associated biological processes. **g)**. An example of the BCAS3 gene with a transcript TSS derived from a human-specific TE insertion. The L1PA2-derived TSS of transcript ENST00000588874.5 is absent in the genomes of chimpanzee, mouse, and chicken, whereas the TSS of the canonical non-TE transcript ENST00000589222.5 is conserved across all four species. The middle sashimi plot shows reads supporting the junction between the L1PA2-derived exon and the second exon in a male esophagus sample. The bottom protein structure plot reveals that the L1PA2-derived transcript lacks the N-terminal region present in the non-TE transcript. **h)**. The expression levels of two transcripts of the BCAS3 gene across human adult tissues.

Compared to transcripts initiated from canonical non-TE promoters, we identified 1166 TE-derived protein-coding transcripts from 1009 protein-coding genes that encode proteins of varying lengths [Figure 4d, supplementary figure 4a, supplementary file 6]. TE-derived protein-coding transcripts contribute to over half of 370 gene expression levels, underscoring their significant impact on transcriptome composition and potential functionality [Figure 4d, supplementary file 6]. We further checked the tissue-specificity of 370 TE-derived protein-coding transcripts that expressed over 50% of total gene expression and produced distinct protein lengths. 83 and 78 transcripts can be identified in all the female and male tissues, respectively, and 77 genes can be found in both sexes [Fig-4d], including several very important genes, such as several components of Nuclear Pore Complex, including NPIPA1/5 and NPIPB2/9/11, and WNT2B. We only identified 5 female-specific protein-coding TE-derived genes, but there are 39 male-specific genes, including 31 testis-specific protein-coding TE-derived genes [Figure 4e].

WNT2B is a member of the wingless-type MMTV integration site (WNT) family and plays an important role in human development as well as carcinogenesis^33,34^. WNT2B produces three distinct transcripts, including one non-TE-derived transcript (ENST00000369684.4) and two TE-derived transcripts (ENST00000256640.9 and ENST00000369686.9), originating from MIR and Arthur1B elements, respectively [Fig-4f, supplementary figure 4b-c]. MIR-derived WNT2B transcript (ENST00000256640.9) only encoded a peptide containing 299 amino acids (AAs), which commonly existed in the other two transcripts. Arthur1B-derived transcript (ENST00000369686.9) can encode an extra 73-AAs at the N-terminal, and the canonical non-TE TSS-derived transcript encodes another 92-AAs N-terminal peptide and results in a 391-AAs protein. As previously reported, these three transcripts differ in protein composition and subcellular localization^33,34^. MIR-derived WNT2B protein is mainly located in the nucleus, and Arthur1B-derived WNT2B protein can be located in both the nucleus and mitochondria. However, non-TE TSS-derived WNT2B protein mainly existed on the endoplasmic reticulum (ER). The TE-derived transcripts are more dominantly expressed across most tissues than the non-TE transcript. In the esophagus (34-year-old male), we observed significantly higher splice junctions from TE-derived 1^st^ exon to their 2^nd^ exon when compared to non-TE TSS-derived WNT2B transcript [Figure 4g]. We further confirmed that the MIR-derived WNT2B transcript was the major expressed isoform in multiple cancer cell lines, including prostate carcinoma Du145, non-small cell lung carcinoma H1299, and triple-negative breast cancer MDA-MB231 [Supplementary Figure 4b,4c]. This highlights the functional significance of TE-derived transcripts in contributing to protein diversity and subcellular dynamics.

### Human-specific TE-derived transcripts

Analysis of evolutionary conservation revealed that the majority of TEs contributing to the transcription of protein-coding and lincRNA genes in the human genome are conserved within primates or mammals. Most of these TE integrations into the human genome occurred within the last 200 million years [Figure 5a, supplementary file 7]. Among mammalian- and primate-conserved TEs, subfamilies such as MIR, MIRb, MIRc, and L2a/b/c contributed the highest number of TE-derived transcription start sites (TSSs). In contrast, human-specific TE subfamilies, including AluYj4, L1HS, and LTR5_Hs, significantly contributed to TE-derived TSSs in the human genome. Comparative genome alignment between humans and three vertebrates—chimpanzees, mice, and chickens—identified 3402 TEs shared between humans and chimpanzees [Figure 5b, supplementary file 8]. Additionally, we identified 68 TEs that are uniquely present in the human genome, of which 50 TEs initiate protein-coding genes, primarily derived from LINE and SINE elements. The remaining 18 TEs, which predominantly belong to LTR elements, initiate lincRNA genes [Figure 5c].

Subfamily-level enrichment analysis revealed distinct activation patterns of TEs in the human genome [Figure 5d]. Notably, LTR12 subfamilies and HERVE-int were strongly enriched among TEs deriving both protein-coding and lincRNA genes. Furthermore, L1PA2 showed unique enrichment as a human-specific active TE subfamily contributing to the transcription of both protein-coding and lincRNA genes. These findings underscore the pivotal role of specific TE subfamilies in shaping the human transcriptome and their evolutionary contributions to transcriptional innovation.

The biological processes related to lipid catabolism and transportation were significantly enriched among genes utilizing Human-Chimpanzee conserved TEs and Human-Chimpanzee-Mouse conserved TEs [Figure 5e, supplementary file 9]. Gene ontology analysis of genes transcribed from 50 human-specific TE-derived transcription start sites (TSSs) revealed enrichment in fatty acid metabolism, purine metabolic processes, chromosome separation, and response to starvation [Figure 5f, supplementary file 9]. Many of these genes are predominantly transcribed from their TE-derived TSSs, including CDT1, TRIB3, ACOT1, and BCAS3. BCAS3 performs multiple critical functions, such as acetyltransferase activation, beta-tubulin binding, and interaction with histone acetyltransferases. It also contributes to cellular responses to estrogen stimulation, enhances catalytic activity, and promotes RNA polymerase II-mediated transcription regulation^35^. An example of human-specific TE activity is one L1PA2 insertion located within the intron between the 9th and 10th exons of BCAS3, generating a novel intragenic TSS. This TSS initiates the truncated transcript ENST00000588874, which encodes a 684-amino-acid (AA) protein. Compared to the canonical full-length protein (928 AA), this truncated version lacks the N-terminal 244 AA [Figure 5g]. This human-specific TE-derived transcript is broadly expressed across all human tissues and has become the dominant transcript in most tissues, with the exception of the artery, colon, and skeletal muscle [Figure 5h]. This highlights the evolutionary significance of TE-derived TSSs in shaping transcriptional landscapes and functional protein diversity in humans.

## Discussion

Transposable elements (TEs) are widespread throughout the human genome and serve as a critical reservoir of regulatory potential, profoundly influencing gene expression and genome evolution. By contributing functional sequences such as enhancers, silencers, and alternative promoters, TEs play a pivotal role in shaping the regulatory landscape of the genome. Although the majority of TEs are silenced in the mammalian genome, increasing evidence reveals that specific TEs actively participate in gene regulation, particularly during early developmental stages. For example, endogenous retroviral elements (ERVs), a subset of TEs, frequently provide enhancer or promoter activity, contributing to the regulation of immune responses, placental development, and early embryogenesis^36-40^. LINE-1 elements are implicated in controlling pluripotency-associated genes during early development^20,41,42^, while LTR elements, such as HERV-H and HERV-K, drive transcriptional activation of critical genes in stem cells^43-45^. Despite these findings, the extent to which TEs influence gene transcription in adult tissues remains less understood. To address this, we systematically analyzed 17,329 human transcriptomes spanning 47 adult tissues to explore the contribution of TEs as gene promoters in regulating transcription. This comprehensive analysis provides crucial insights into the role of TEs in expanding regulatory complexity and shaping the human transcriptomic landscape across diverse biological contexts.

Our findings reveal that TE-derived transcripts are broadly expressed in human tissues, playing dual roles in maintaining housekeeping functions and driving tissue-specific gene regulation. We identified that 21% of human genes, comprising 5,330 protein-coding and 2,673 non-coding genes, utilize TE-derived transcription start sites (TSSs) to transcribe alternative isoforms in different tissues. Many of these isoforms are lowly expressed, particularly TE-derived lincRNAs. Notably, approximately 30% of these transcripts (1,300 in females and 1,619 in males) dominantly contribute to over 50% of their host gene expression in at least one tissue, demonstrating their significant functional impact. In males, 401 TE-derived transcripts are tissue-specific, with 319 exclusively expressed in the testis, highlighting their critical role in the unique epigenetic and transcriptomic landscape of male germ-line cells. Among TE-derived TSSs, SINE elements accounted for nearly half of those associated with protein-coding genes, while LTR elements contributed about half of the TSSs for lincRNA genes, consistent with previous findings that LTRs are a major source of lincRNAs in the mammalian genome^46,47^. Cross-tissue comparisons revealed strong associations between TE-derived transcripts and unique tissue functions, suggesting that TEs significantly contribute to tissue specificity and are likely regulated by tissue-specific transcription factors. These findings underscore the prevalence of TE-derived transcription and suggest that TEs have been domesticated as alternative promoters, enhancing tissue-specific gene regulation during evolution. This highlights their pivotal role in shaping the human transcriptomic landscape and expanding regulatory complexity.

We observed pronounced sex-specific expression of TE-derived transcripts regulated by sex hormones, particularly in breast tissue. In this tissue, 312 protein-coding and 83 lincRNA transcripts exhibited strong sex-biased expression, with many being highly expressed in females and associated with mammary gland development. TE subfamily analysis revealed that MIRb and L2b elements contributed to sex-biased protein-coding transcripts, while HERV15-int and LTR12C were associated with lincRNAs. Female-biased TEs included L2a and AluJb, whereas AluY exhibited male-specific activation. Nearly half of these sex-biased TE-TSSs harbor sex hormone receptor binding sites, such as ESR1 and AR, consistent with previous studies showing TEs as rich reservoirs of ER and AR binding sites in the human genome, which can be aberrantly activated in cancer cells. Our findings further suggest that TEs containing ER or AR binding sites actively contribute to normal gene regulation in human sex organs, underscoring their role in both physiological and pathological processes.

We demonstrated that TE-derived alternative transcription initiation plays a significant role in enhancing protein translation diversity. Approximately 40% of TE-derived transcripts in our study encode protein products, while nearly 60% are non-translatable, such as retained-intron or processed transcripts. TE-derived protein-coding transcripts were shown to influence key biological pathways, including inflammatory responses, PLC signaling, and TGF signaling. Among 370 TE-derived protein-coding transcripts contributing over 50% of gene expression, distinct tissue-specific and sex-specific expression patterns were evident. Notably, TE-derived protein-coding transcripts can introduce novel N-terminal peptides, which critically influence subcellular localization, as exemplified by WNT2B. These findings underscore the functional and evolutionary significance of TEs in expanding the transcriptomic and proteomic landscape of humans. Additionally, we identified 68 human-specific TE-derived transcripts linked to essential biological processes such as metabolism and environmental adaptation. These human-specific transcripts highlight the unique role of TEs in shaping species-specific traits and enabling adaptation to environmental pressures, providing a distinct evolutionary advantage that sets human evolution apart from other primates.

Together, our findings highlight the pivotal role of transposable elements (TEs) in human gene regulation. By serving as promoters, TE-derived transcription start sites (TE-TSSs) play a crucial role in driving transcriptional and translational innovation within the human genome. These results provide valuable insights into how TEs enhance the regulatory and functional complexity of the human transcriptome, enabling adaptation and fostering evolutionary innovation as dynamic and novel genomic elements.

## Method

### Transposable element-derived transcripts expressed in human adult tissues

The transposable element (TE) data for the human genome (hg38) was obtained from the UCSC database (https://hgdownload.soe.ucsc.edu/goldenPath/hg38/database/rmsk.txt.gz)^48^. Simple repeats and satellite elements were excluded from this analysis, focusing only on TEs belonging to the DNA, LINE, SINE, and LTR classes.

Gene and transcript annotations for the human genome (version 26) were downloaded from the GENCODE database (https://www.gencodegenes.org/human/release_26.html)^49^. From these annotations, the genomic coordinates of transcription start sites (TSSs) were extracted. To identify transcripts with TEs-derived TSSs, the intersectBed function from the Bedtools suite was used^50^. Transcripts whose TSSs overlapped with TE regions were classified as TE-derived transcripts.

The read count table of transcripts from 43 male and 44 female human adult tissues was obtained from the GTEx Analysis V8 dataset based on GENCODE v26 annotations (https://gtexportal.org/home/downloads/adult-gtex/bulk_tissue_expression), excluding cell line samples^51^. The data were separated into male and female tissue datasets, and transcripts with low expression (CPM ≤ 1 in at least half of the samples in a tissue) were filtered out separately for each sex. After filtering, a total of 102,082 expressed transcripts were identified, including 92,612 derived from protein-coding and lincRNA genes. The Trimmed Mean of M-values (TMM) method was applied to normalize the read counts across all samples, and RPKM values were calculated for individual transcripts in each sample. Additionally, the average RPKM of each transcript was computed separately for male and female tissues. TE-derived transcripts were further filtered, retaining those with an average RPKM ≥ 0.1 in at least one tissue group, resulting in 5,850 TE-derived transcripts from 3,528 protein-coding and 944 lincRNA genes. Z-scores of these transcripts’ expression levels were calculated across male and female tissues and used for clustering via the k-means method in R^52^. Functional enrichment analysis was performed using the clusterProfiler package^53^, revealing biological pathways and functions associated with genes harboring TE-derived transcripts in different clusters. This comprehensive analysis highlights the role of TE-derived transcripts in human adult tissues.

### Subfamily Enrichment of TEs that derived TSS of transcripts

To assess the enrichment of TE subfamilies contributing to transcripts derived from protein-coding and lincRNA genes, the Log Enrichment Ratio (LER) was calculated. This was determined by comparing the fraction of a given TE subfamily among all TEs associated with expressed transcripts (TE-derived TSSs) to the fraction of the same subfamily within the total TEs in the human genome, using the following formula:

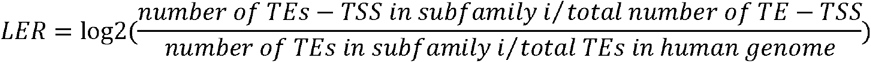

The individual copy numbers for each TE subfamily were counted using the TE annotation dataset described earlier. To identify enriched TE subfamilies contributing to transcripts, the data were filtered based on two criteria: (1) the number of individual TE copies in a subfamily must exceed 5, and (2) the Log Enrichment Ratio (LER) value must be greater than 2.

### Differential expressed TE derived transcripts between female and male in tissues

The expression levels of transcripts from individual samples within each tissue were analyzed to identify differentially expressed TE-derived transcripts (sex-DET) between females and males. The difference in expression (FDR) for each TE-derived transcript was calculated using the wilcox.test method, with adjustments made for multiple testing using the p.adjust function in R with the FDR method. Fold changes in expression were determined by dividing the average expression of each TE-derived transcript in males by the average expression in females. Sex-DETs within tissues were identified based on the following criteria: (1) FDR < 0.01, (2) absolute log2 fold change > log2(1.5), (3) average expression ≥ 0.1 in either the female or male group, and (4) exclusion of genes located on the X and Y chromosomes. Additionally, sexual tissues (testis, prostate, ovary, vagina, and uterus) were excluded from this analysis. The ggplot2 and ggrepel packages in R were used to generate volcano plots to visualize sex-DETs for both protein-coding and lincRNA transcripts^54,55^. Enrichment analysis of the identified sex-DETs was conducted using the ToppFun tool (https://toppgene.cchmc.org/enrichment.jsp)^56^. This approach highlights the biological significance of TE-derived transcripts with differential expression patterns between sexes.

Binding motif regions for the ESR1 and AR transcription factors in the human genome were obtained from the ReMap2022 database (https://remap.univ-amu.fr/about_hsap_page)^57^. The intersectBed function was then used to determine whether ESR1 and AR binding motifs overlapped with transposable elements (TEs) or their 500 bp flanking regions. The bigWig file for ESR1 ChIP-seq data from the human Ishikawa cell line treated with 10 nM estradiol for 1 hour was downloaded from ENCODE (https://www.encodeproject.org/experiments/ENCSR000BIY/)^58,59^. Similarly, bigWig and bam files of total RNA-seq data for female adult human breast epithelium tissue (51 years) and male adult human breast epithelium tissue (37 years) were retrieved from ENCODE (ENCSR438YPF and ENCSR094VRQ, respectively). Visualization of ChIP-seq and RNA-seq tracks was performed using the WashU Epigenome Browser (https://epigenomegateway.wustl.edu/) with the bigWig files^60^. The bam files of RNA-seq data were analyzed using IGV (v2.17.0) to generate sashimi plots and to count the number of reads supporting exon junctions^61^. Those tracks of WashU Epigenome browser and IGV can be accessed through the GitHub (https://github.com/BenpengMiao/TE_transctipts_in_human_adult_tissues/tree/main/Track). This workflow enabled a detailed examination of TE-associated regulatory motifs and transcriptional activity in sex-specific contexts.

### Expression contribution of TE-derived transcripts in the genes

For individual samples across all tissues, the gene expression was calculated by the sum of all the expressed transcript from the same gene, and then the expression contribution of TE-derived transcript in the gene was measured by the following formula:

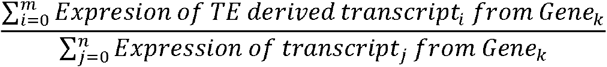

The *m* is the total TE-derived transcripts in the gene; the *n* is the total transcripts in the same gene, including the TE-derived transcripts and other non-TE-derived transcripts.

The average expression contribution of TE-derived transcripts was calculated separately for each tissue group, male and female. Simultaneously, the mean expression level of each gene was determined within each tissue group. If a gene’s mean expression was less than 0.1, the expression contribution of its TE-derived transcripts was set to zero.

### TE-derived protein-coding transcripts

Gene types and transcript biotypes were extracted from the GTF annotation file of GENCODE v26 (https://ftp.ebi.ac.uk/pub/databases/gencode/Gencode_human/release_26/gencode.v26.annotation.gtf.gz). Protein-coding transcript translation sequences were also retrieved from GENCODE v26 (https://ftp.ebi.ac.uk/pub/databases/gencode/Gencode_human/release_26/gencode.v26.pc_translations.fa.gz) and used to calculate peptide lengths. This analysis aimed to determine whether TE-derived transcripts produce protein products with structural differences compared to non-TE-derived transcripts from the same gene. A total of 1,759 protein-coding transcripts were identified as TE-derived, with 1,116 of them producing proteins that differed structurally from those encoded by non-TE-derived transcripts.

Exon structures and protein products of TE- and non-TE-derived transcripts from the same gene were obtained from Ensembl (https://useast.ensembl.org/index.html) and UniProt (https://www.uniprot.org/)^62,63^. Structural differences in protein products were further analyzed using the AlphaFold Protein Structure Database (https://alphafold.ebi.ac.uk/)^64,65^. Protein structures were visualized and modified using the Structure Viewer available in this database.

For RNA-seq analysis, the BAM file of polyA+ RNA-seq data from male adult human esophagus tissue (34 years old) was downloaded from ENCODE (https://www.encodeproject.org/experiments/ENCSR102TQN/). This BAM file was analyzed using IGV (v2.17.0) to generate sashimi plots and quantify the number of reads supporting junctions between TE-derived first exons and subsequent exons and the tracks are available in GtiHub (https://github.com/BenpengMiao/TE_transctipts_in_human_adult_tissues/tree/main/Track). This analysis provided insights into the transcriptional and structural impacts of TE-derived transcripts.

### RT-qPCR

Total RNA was extracted using Monarch Total RNA Miniprep Kit (NEB #T2010) from cultured cancer cell lines Du145, H1299 and MDA-MB-231. cDNA was generated from 500ng total BioRad RNA using iScript™ Reverse Transcription Supermix for RT-qPCR. Quantitative RCR was performed using PowerUp™ SYBR™ Green Master Mix for qPCR with the primers listed in the following. GAGPH was used as the endogenous reference to normalize the expression of the targets.

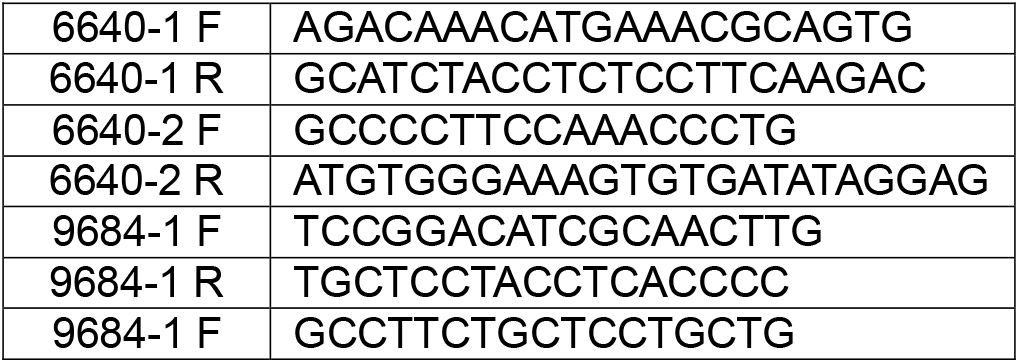
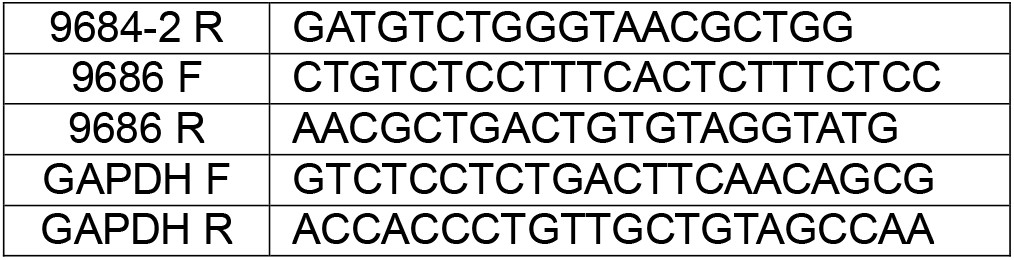

### Evolutionary conservation of TEs that derived transcripts

The taxonomic information for TE subfamilies was retrieved from the Dfam database (https://dfam.org/home)^66^, and their estimated evolutionary ages (in million years) were calculated using the TimeTree 5 database (https://timetree.org/) based on taxonomic classifications^67^. A total of 26 distinct taxa were identified for subfamilies that generate transcripts, which were grouped into four major taxonomic categories: Homo, Primate, Mammalia, and Amniota, based on evolutionary timeframes. The TEs contributing to transcripts in the human genome were aligned to the chimpanzee, mouse, and chicken genomes using the liftOver tool with default parameters (https://genome.ucsc.edu/cgi-bin/hgLiftOver)^68^. TEs successfully mapped to the other genomes were classified as orthologous, and based on their orthologous patterns, they were divided into four evolutionary groups: orthologous in all three genomes (chimpanzee, mouse, and chicken; H-C-M-C), orthologous in chimpanzee and mouse genomes (H-C-M), orthologous only in the chimpanzee genome (H-C), and specific to the human genome (H). Subfamily enrichment analysis for TEs in these evolutionary groups was performed using previously described methods. Gene enrichment analysis for transcripts derived from TEs in each group was conducted using the clusterProfiler package in R, with enriched biological processes and associated genes visualized using the emapplot and cnetplot functions. This integrative analysis highlights the evolutionary dynamics of TEs and their functional contributions to human transcriptional regulation.

## CODE AVAILABILITY

The source code, documentation, and supporting data of this work are freely available at https://github.com/BenpengMiao/TE_transctipts_in_human_adult_tissues.

## AUTHOR CONTRIBUTIONS

T.P.W. and B.A.Z. conceived and designed the study, and developed the methodology. B.M., A.A., and Y.Y. performed the computational analysis and data visualization. X.L. performed experiments. B.M., T.P.W. and B.A.Z. contributed to the manuscript writing.

## ACKNOWLEDGEMENTS

The National Institutes of Health supported this work: R35GM142917(BZ), U24HG012070(BZ), U24NS132103(BM). Cancer Prevention Research Institute of Texas supported this work: CPRIT-RR180072 (TW). Funding for open access charge: National Institutes of Health.

## Abbreviations

SB: Adipose: Subcutaneous
VO: Visceral Omentum
Adrenal_GA: Adrenal Gland
Breast_MT: Breast Mammary Tissue
Kidney_CRx: Kidney_Cortex
AR: Artery: Aorta
CR: Coronary
TB: Tibial
AY: Brain: Amygdala
ACC: Anterior_cingulate_cortex_BA24
CR: Cortex
CH: Cerebellar_Hemisphere
CBG: Caudate_basal_ganglia
FC: Frontal_Cortex_BA9
HIP: Hippocampus
HYP: Hypothalamus
NAC: Nucleus_accumbens_basal_ganglia
PUT: Putamen_basal_ganglia
SCC: Spinal_cord_cervical_c_1
SN: Substantia_nigra
SG: Colon: Sigmoid
TA: Transverse
GJ: Esophagus: Gastroesophageal_Junction
MC: Mucosa
MS: Muscularis
AA: Heart: Atrial_Appendage
LV: Left_Ventricle
Minor_SG: Minor Salivary Gland
Not_Sun_SP: Skin: Not_Sun_Exposed_Suprapubic
Skin_Sun_LL: Sun_Exposed_Lower_leg

