## supplementary_materials for "Expression spectrum of TE-derived transcripts in human adult tissues"

**Supplementary figure 1. TE-derived transcripts expressed in human adult tissues.**

**Supplementary figure 2. The subfamily distribution of TEs contributing to differentially expressed transcripts between females and males (sex-DETs) in breast tissue.**

**Supplementary figure 3. The classes of TEs contributing to transcripts from protein-coding genes and lincRNAs in females and males.**

**Supplementary figure 4. TE-derived protein-coding transcripts with altered protein length.**

**supplementary file 1. Expression of TE-derived transcript in human adult tissues.**

**supplementary file 2. Z-score Expression cluster transcript.**

**supplementary file 3. sex differential DET in tissues.**

**supplementary file 4. Expression contribution of TE-derived transcripts in genes across tissues.**

**supplementary file 5. Enriched BP based on different transcript types.**

**supplementary file 6. TE-derived protein-coding transcripts.**

**supplementary file 7. Dfam Taxa of TEs that derived transcripts.**

**supplementary file 8. LiftOver orthologous of TEs that derived transcripts.**

**supplementary file 9. GO terms for different evolutionary groups.**

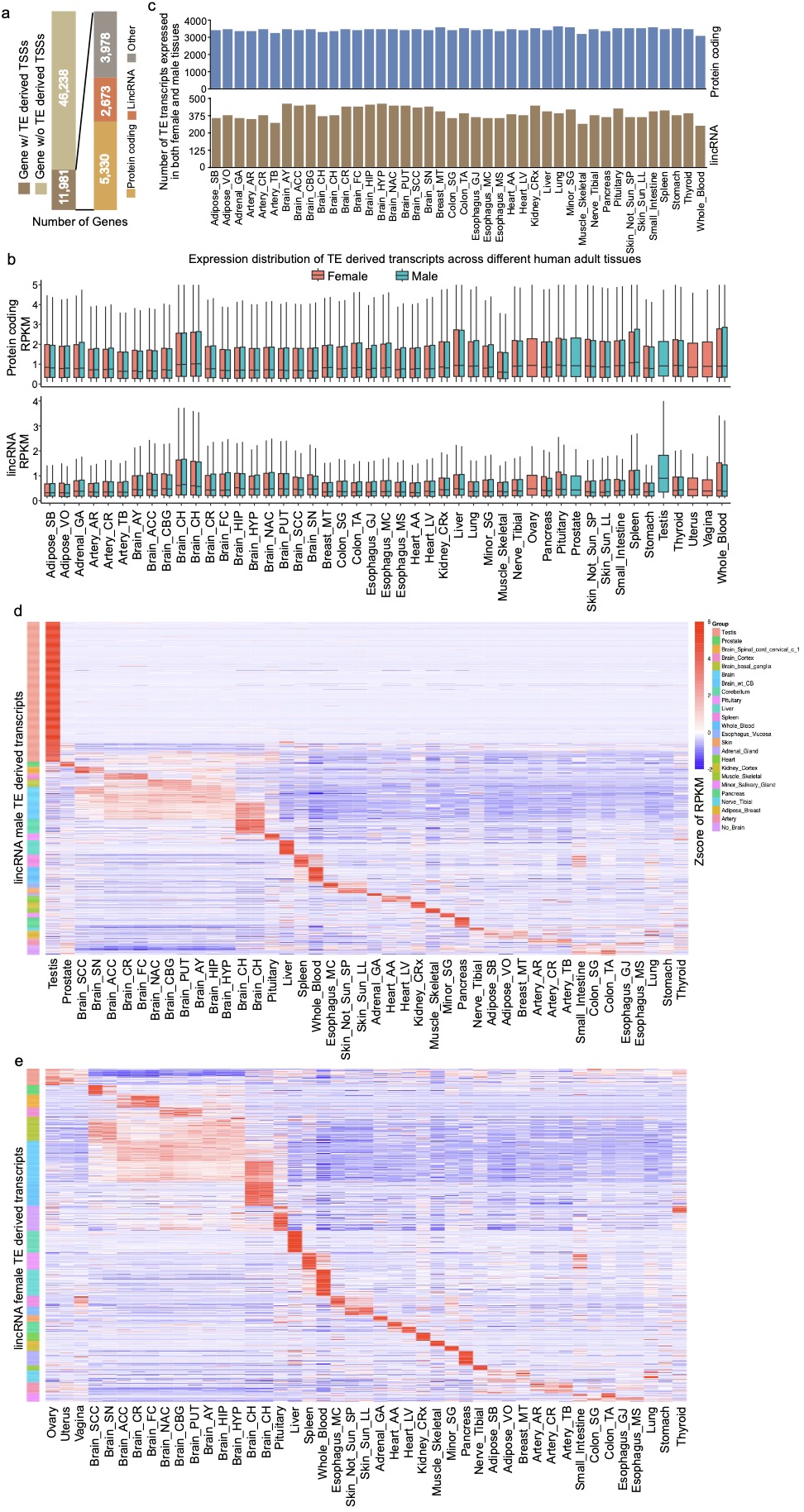

**Supplementary figure 1. TE-derived transcripts expressed in human adult tissues.**

**a)** The number of genes containing TE-derived transcripts based on GENCODE v26 annotation. A total of 5,330 protein-coding genes and 2,673 lincRNAs have TE-derived transcripts. **b).** A boxplot showing the expression distribution of TE-derived transcripts from protein-coding genes and lincRNAs across tissues. Red and blue colors represent female and male samples, respectively. **c).** The number of TE-derived transcripts from protein-coding genes and lincRNAs expressed in both female and male tissues. Over 3,000 protein-coding and more than 300 lincRNA TE-derived transcripts were identified in both sexes. **d-e).** Z-scores of lincRNA TE-derived transcript expression for males and females, respectively. Different colors indicate distinct tissue groups.

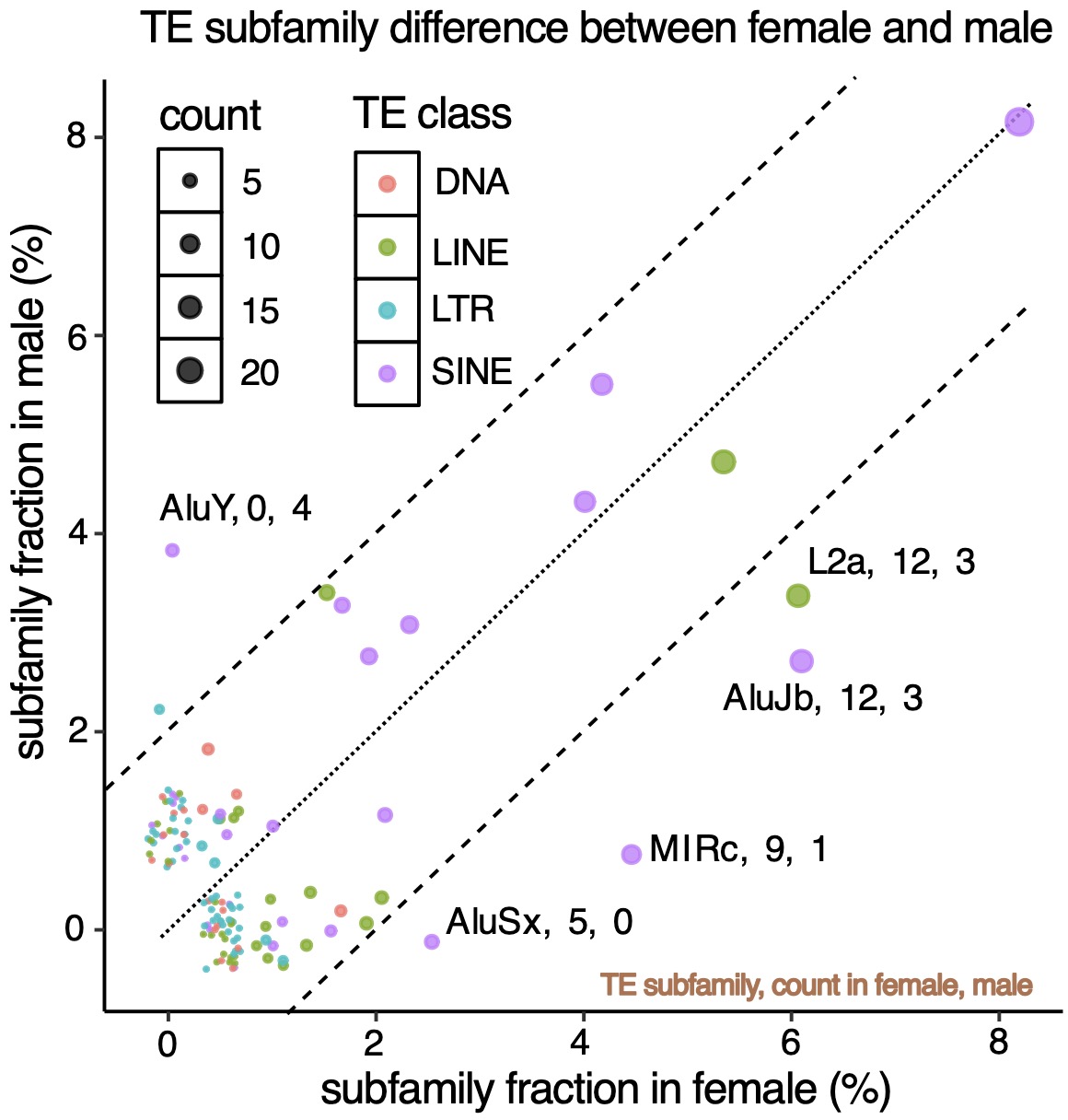

**Supplementary figure 2. The subfamily distribution of TEs contributing to differentially expressed transcripts between females and males (sex-DETs) in breast tissue.** Each dot represents a TE subfamily, with dot size indicating the number of TEs within the subfamily that derived sex-DETs in breast tissue. Colors represent different TE classes. The x-axis shows the fraction of a subfamily contributing to up-regulated sex-DETs in females, while the y-axis shows the fraction contributing to up-regulated sex-DETs in males. For example, 12 TEs from the L2a subfamily derived up-regulated sex-DETs in females, while only 3 contributed to males. Conversely, 4 AluY elements derived up-regulated sex-DETs in males, but none contributed to females.

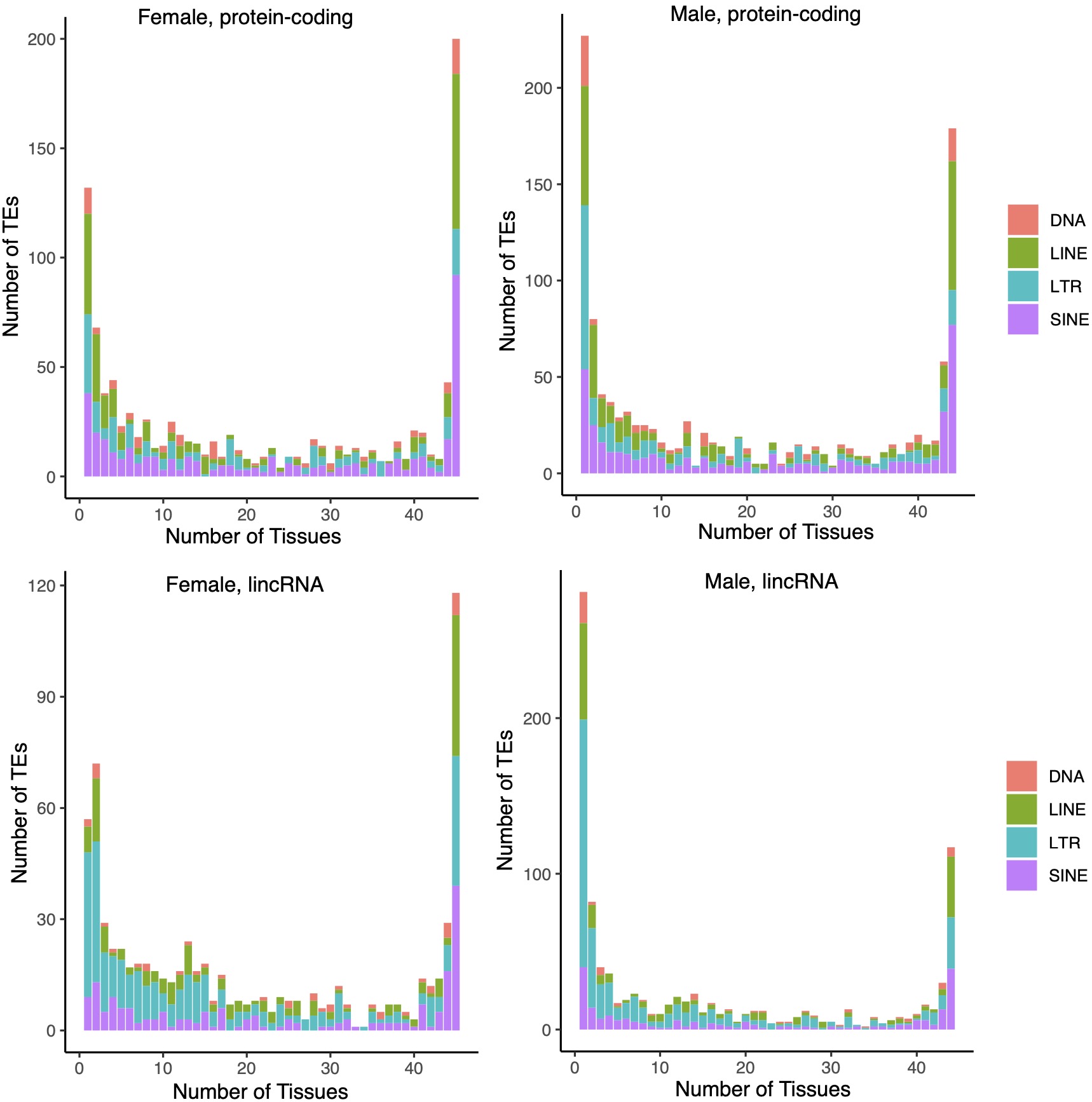

**Supplementary figure 3. The classes of TEs contributing to transcripts from protein-coding genes and lincRNAs in females and males.** The x-axis represents the number of tissues sharing the same expressed TE-derived transcripts, while the y-axis indicates the number of TEs. Bar colors represent different TE classes.

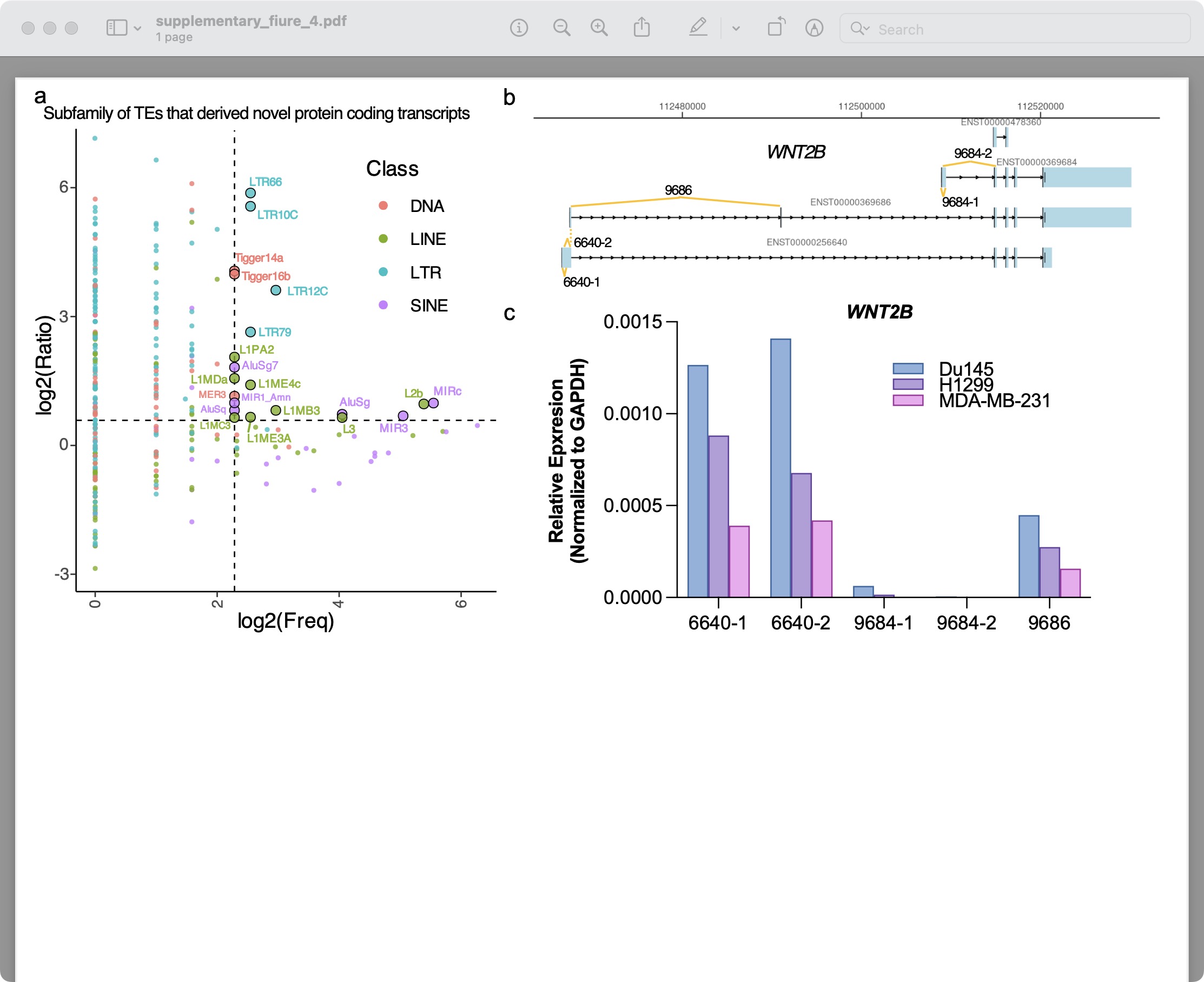

**Supplementary figure 4. TE-derived protein-coding transcripts with altered protein length.**

**a).** Subfamily enrichment of TEs generating protein-coding transcripts with protein lengths differing from canonical non-TE-derived transcripts. The x-axis represents the number of individual TEs within a subfamily, while the y-axis shows the log2 ratio of the fraction of subfamily-derived protein-coding transcripts with altered protein length to the fraction of the subfamily in the total TEs of the human genome. Different colors indicate TE classes. **b).** A schematic diagram of all WNT2B transcripts retrieved from GRCh38.p13 (Ensembl v109). Yellow lines denote the locations where RT-qPCR primers were designed. **c)**. RT-qPCR analysis showing the relative expression of WNT2B transcripts (normalized to GAPDH) in three cancer cell lines: Du145 (prostate cancer), H1299 (lung cancer), and MDA-MB-231 (breast cancer). 6640-1/2 represents the MIR-derived transcript (ENST00000256640.9), 9686 represents the Arthur1B-derived transcript (ENST00000369686.9), and 9684-1/2 represents the canonical transcripts (ENST00000369684.4). Bar colors correspond to the three cell lines
